# Nosocomial *Pseudomonas aeruginosa* regulates alginate biosynthesis and Type VI secretion system during adaptive and convergent evolution for coinfection in critically ill COVID-19 patients

**DOI:** 10.1101/2021.04.09.439260

**Authors:** Jiuxin Qu, Zhao Cai, Xiangke Duan, Han Zhang, Shuhong Han, Kaiwei Yu, Zhaofang Jiang, Yingdan Zhang, Yang Liu, Yingxia Liu, Lei Liu, Liang Yang

**Author notes:** Corresponding authors: Lei Liu and Liang Yang. These authors contributed equally to this work (Jiuxin Qu, Zhao Cai). These authors contributed equally to this work and share the corresponding authorship (Lei Liu, Liang Yang).

## Abstract

COVID-19 pandemic has caused millions of death globally and caused huge impact on the health of infected patients. Shift in the lung microbial ecology upon such viral infection often worsens the disease and increases host susceptibility to secondary infections. Recent studies have indicated that bacterial coinfection is an unignorable factor contributing to the aggravation of COVID-19 and posing great challenge to clinical treatments. However, there is still a lack of in-depth investigation on the coinfecting bacteria in COVID-19 patients for better treatment of bacterial coinfection. With the knowledge that *Pseudomonas aeruginosa* is one of the top coinfecting pathogens, we analyzed the adaptation and convergent evolution of nosocomial *P. aeruginosa* isolated from two critical COVID-19 patients in this study. We sequenced and compared the genomes and transcriptomes of *P. aeruginosa* isolates longitudinally and parallelly for its evolutionary traits. *P. aeruginosa* overexpressed alginate and attenuated Type VI secretion system (T6SS) during coinfection for excessive biofilm formation and suppressed virulence. Results of bacterial competition assay and macrophage cytotoxicity test indicated that *P. aeruginosa* reduced its virulence towards both prokaryotic competitors and eukaryotic host through inhibiting its T6SS during evolution. *P. aeuginosa* T6SS is thus one of the reasons for its advantage to cause coinfection in COVID-19 patients while the attenuation of T6SS could cause a shift in the microecological composition in the lung. Our study will contribute to the development of therapeutic measures and the discovery of novel drug target to eliminate *P. aeruginosa* coinfection in COVID-19 patient.

## Introduction

Coronavirus Disease 2019 (COVID-19) induced by Severe Acute Respiratory Syndrome Coronavirus 2 (SARS-CoV-2) has caused over 2.8 million death globally as reported by WHO at Apr 2021. SARS-CoV-2 induces cytokine storm and hyperinflammation resulting in respiratory dysfunction and necrosis of epithelial cells of the lungs [1, 2]. Such histological changes in the lung tissues may result in dysbiosis of lung microbiome and give rise to secondary bacterial infection in COVID-19 patients [3]. Several groups of researchers have emphasized that microbial coinfection indeed occurs in COVID-19 patients, especially in severe cases, exacerbates disease progress and makes difficulties to clinical treatments [4-7].

Nosocomial bacterial pathogens are risk factors contributing to such microbial coinfections and resulting in increased death of critically ill COVID-19 patients [8, 9]. Bacterial coinfection in COVID-19 patients is a serious problem which should not be neglected. The common coinfecting bacteria reported include *Mycoplasma pneumonia, Staphylococcus aureus, Klebsiella spp*., *Acinetobacter baumannii, Haemophilus influenzae, Pseudomonas aeruginosa* and etc [10, 11]. There was bacteremia case caused by *Pseudomonas* reported [12]. *P. aeruginosa* is identified as a top coinfecting bacteria in our hospital (unpublished data).

*Pseudomonas aeruginosa* is a well-known nosocomial pathogen causing fatal chronic infections in immunocompromised patients with diseases like cystic fibrosis, catheter-associated infections and burn wounds [13, 14]. This bacterium induces infections through different mechanisms, including biofilm formation, quorum sensing systems and secretion of virulence factors [13]. *P. aeruginosa* possesses diverse virulence systems including quorum sensing systems, protein secretion systems, lipopolysaccharides etc for competition in the polymicrobial environments and invasion to host cells[13]. Longitudinal analysis of convergent evolution of *P. aeruginosa* clinical isolates indicated that remodeling of virulence secretion is essential for host adaptation [15, 16]. Mutations in genes, *mucA, vgrG, lasR, rpoN* and *pvdS*, significantly impair quorum sensing systems, protein secretion systems and virulence factor biosynthesis allowing *P. aeruginosa* to escape from host immune clearance in patients with lung diseases such as cystic fibrosis and ventilator-associated pneumonia [15-18]. Among the secretion systems, Type VI secretion system (T6SS) highly correlates to bacterial competition and host interaction of *P. aeruginosa* with other organisms [19]. Analysis of these evolutionary traits could contribute to the development of therapeutic measures to treat infections caused by *P. aeruginosa*. However, there is still a paucity of data on the insight of nosocomial *P. aeruginosa* coinfection with SARS-CoV-2 virus and its evolution during coinfection. To address this, we sequenced and compared the genomes and transcriptomes of four *P. aeruginosa* isolates sampled longitudinally from two critically ill COVID-19 patients to investigate its adaptive convergent evolution. The isolates collected from different patients are of the same sequence typing indicating the occurrence of hospital-acquired *P. aeruginosa* infection. The results showed that these isolates form excessive biofilm by developing mucoid phenotype through increasing alginate biosynthesis for long-term colonization in the host. More notably, we demonstrated here that *P. aeruginosa* evolves and attenuates its T6SS, especially HSI-II T6SS, to suppress its virulence to escape host clearance during coinfection. Understanding such adaptive convergent evolution could contribute to the treatment decision and the discovery of novel drug target to eliminate *P. aeruginosa* infection in COVID-19 patients.

## Material and Methods

### Isolate collection, bacterial strains and growth media

Four isolates of *Pseudomonas aeruginosa* were collected at two different time points longitudinally from sputum samples or bronchioalveolar lavage fluids (BALF) of two critically ill COVID-19 patients respectively during routine clinical tests. Isolates collected from patient 1 were named as LYSZa2 and LYSZa3 while isolates collected from patient 2 were named as LYSZa5 and LYSZa6. Luria-Bertani (LB) broth and ABTGC medium supplemented with 10% Tryptic Soy Broth (TSB) were used for growing cultures. ABTGC medium consists of 0.1% MgCl_2_, 0.1% CaCl_2_, 0.1% FeCl_3,_ 0.2% glucose, 0.2% casamino acids and 10% A10 medium which is made of 15.1mM (NH_4_)_2_SO_4_, 33.7mM Na_2_HPO_4_•2H_2_O, 22 mM KH_2_PO_4_ and 0.05mM NaCl.

### Ethical statement

This work does not include any direct participation of the patients and does not contain any identifiable human data. This work is approved by the Ethics Committee of Shenzhen Third People’s Hospital, Second Hospital Affiliated to Southern University of Science and Technology [2020-184] and filed with the Ethics Committee of Southern University of Science and Technology [20200069].

### Antimicrobial susceptibility tests

Antimicrobial susceptibility of the isolates to various antibiotics, including ceftazidime, piperacillin, cefoperazone/sulbactam, imipenem, aztreonam, and levofloxacin, were performed using the Kirby-Bauer disc-diffusion method. Diameters of inhibition zone was measured using a vernier caliper. Susceptibility was determined according to CLSI 2019[20]. Results of antibiotic susceptibility tests were listed in Table S1.

### Biofilm formation assay

The isolates were cultured in LB broth at 37°C for overnight. The overnight cultures were diluted to OD_600nm_ 0.01 in fresh LB broth. 100µL of the diluted cultures were loaded into 96-well plate in triplicates and incubated for 24h at 37°C statically allowing the formation of biofilm. After removing spent media, biofilms were washed carefully with ddH_2_O for two times. Biofilms were then stained by 125 µL of 0.1% crystal violet (CV) with 15 min incubation at room temperature. CV stain in the wells was discarded while stained biofilms were washed twice thoroughly with ddH_2_O and air-dried. Biofilms were then dissolved into 125 µL of 30% acetic acid and quantified relatively by measuring OD_550nm_ values on a Tecan infinity pro200 microplate reader.

### Genome extraction and sequencing

The isolates were cultured in LB broth at 37°C to early stationary phase. For Illumina sequencing, genomic DNA of each isolates was extracted using AxyPerp Bacterial Genomic DNA Miniprep Kit (Corning, New York, USA) following manufacturer’s protocol. PCR-free libraries were constructed using VAHTSTM PCR-Free DNA Library Prep Kit for Illumina® (Vazyme, China) following standard protocol. VAHTSTM DNA Adapters for Illumina® (Vazyme, China) was used to tag adaptor to purified fragments. Quality of the libraries were assessed by Agilent Technologies 2100 Bioanalyzer and qPCR. Paired-end DNA sequencing was then performed on Illumina HiSeq X platform with read length of 150 bp. For PacBio sequencing, genomes of LYSZa2 and LYSZa5 were extracted using Mabio Bacterial DNA Extraction Mini Kits (Mabio) according to manufacturer’s protocol. DNA fragmentation was done using G-tubes (Covaris). SMRTbell DNA template libraries were prepared according to the manufacturer’s specification (PacBio, Menlo Park, USA). DNA sequencing was then performed on Pacific Biosciences RSII sequencer (PacBio, Menlo Park, USA).

### Transcriptome extraction and sequencing

The isolates were cultured in triplicates in LB broth at 37°C to early stationary phase. Magen HiPure Universal RNA Mini kits (MCBio, China) was used to extract total RNA following the manufacturer’s protocol. Extracted RNA was quantified using Qubit 2.0 (Thermo Fisher Scientific, MA, USA) and Nanodrop One (Thermo Fisher Scientific, MA, USA). Quality of RNA samples was assessed by Agilent 2100 system (Agilent Technologies, Waldbron, Germany). RNA libraries were constructed according to standard protocol using NEB Next® Ultra™ Directional RNA Library Prep Kit for Illumina® (New England Biolabs, MA, USA). Ribosomal RNA depletion was carried out using Ribo-zero rRNA Removal Kit. cDNA was synthesized using NEB Next First Strand Synthesis Reaction Buffer. Paired-end RNA sequencing was performed on Illumina NovaSeq 6000 platform with read length of 150 bp.

### Sequencing data analysis

Genomic Illumina sequencing reads were assembled into contigs using De Novo Assembly module of CLC Genomics Workbench 20 (Qiagen) with default parameters. Single nucleotide polymorphism was detected using ‘Resequencing’ module of CLC Genomics Workbench 20 based on frequency of more than 80% using *P*.*aeruginosa* PAO1 genome as reference. PacBio sequencing reads were assembled into draft genome using HGAP4 pipeline of SMRT Link software v9.0 with default settings. Rearrangements of draft genomes were checked using Mauve software v.2.4.0 by PROGRESSIVEMAUVE alignment mode[21]. Multilocus sequence typing (MLST) was performed using MLST service available on the Center of Genomic of Epidermiology (CGE) webserver[22]. Identification of antimicrobial resistance genes was performed using ResFinder service on CGE webserver based on 85% of identity and 60% of minimal length[23]. Phylogenetic tree was constructed using Parsnp under libMUSCLE aligment mode using draft genomes of the isolates and other strains [24]. Genomic islands on LYSZa2 and LYSZa5 genomes were predicted using IslandViewer 4 webservice [25]. Circular plot was constructed using BLAST Ring Image Generator [26]. Genomic loci were visualized using Easyfig package[27]. Gene names on the genomic loci were obtained from Pseudomonas Genome Database[28]. Prediction of protein functions of the genes were performed using NCBI Prokaryotic Genome Annotation Pipeline (PGAP) [29].

RNA sequences were pre-processes, mapped to *P. aeruginosa* PAO1 reference genome and analyzed using RNA analysis module of CLC Genomics Workbench 20 (Qiagen) with default parameters. Total read counts of each sample were normalized and compared using DESeq2 R package. Differentially expressed genes (DEGs) were selected based on absolute fold change≧4, adjusted p-value<0.05 and base mean ≧ 20 [30]. GO enrichment analysis of DEGs was performed on DAVID bioinformatics database v6.8 [31]. PCoA plot and heatmap were drawn using Vegan, ggplot2, and pheatmap packages in R 4.0.0.

### Bacterial competition assay

Competition between *P. aeruginosa* and *Escherichia coli* was carried out following the steps described previously by Hachani.et.al[32]. Briefly, the isolates were cultured overnight at 37°C on LB agar plates. *E. coli/pLacZ* strain was grown on LB agar plates containing 40 mg/mL of 5-bromo-4-chloro-indolyl-β-D-galactopyranoside (X-gal) at 37°C for overnight and appeared blue in colour. Single colony of each strain was subcultured in TSB medium and grown under agitation for overnight. Appropriate volume of each overnight culture was taken to make final OD600nm =1 and centrifuged to collect cells. Cell pellets were resuspended in 100 µL TSB medium while 10 µL of each suspension was spotted and incubated on LB agar plate for 5 hours at 37°C. 30 µL of cell suspension of each *P. aeruginosa* strain was taken and mixed gently with 30 µL of *E. coli* cell suspensions. 20 µL of mixed cell cultures were spotted onto LB agar plate and incubated for 5 hours at 37°C. Each bacterial spot was scraped from LB agar plates and resuspended in 1mL of TSB medium. Resuspensions of bacterial spots were diluted to 10^−3^ serially in 10-fold dilutions. These serial cell dilutions were then spotted in triplicates onto LB agar plates containing 40 mg/mL X-gal and incubated for 8 hours for killing *E. coli*. 100 µL of 10^−3^ dilution of each mixed bacterial spot were plated onto LB agar plates containing 40 mg/mL X-gal for CFU counting of *E. coli* for quantitating killing effect.

### Cytotoxicity assay

The cytotoxicity of the isolates was assayed by using murine RAW 264.7 macrophages. RAW macrophages were grown in 24-well plates in Dulbecco modified Eagle medium (DMEM), GlutaMAX, sodium pyruvate, and phenol red supplemented with 10% FBS. Prior to infection, confluent RAW cells were washed twice with sterile PBS and incubated in DMEM medium. Log-phase cultures were washed with sterile PBS twice, and resuspended in DMEM medium devoid of FBS. Macrophage cells were infected with the isolates respectively at a Multiplicity of Infection (MOI) of 20 at 37 °C in 5% CO2 incubator. After 3 h infection, the culture supernatants were collected for detecting lactate dehydrogenase (LDH) activities. The LDH activities were detected by using commercially LDH cytotoxicity kit (YAESEN Bio) according to standard procedure.

### Data Availability

All Illumina sequencing data used in this study could be found from BioProject No. PRJNA706783 and assembled genomes of *P. aeruginosa* LYSZa2 and *P. aeruginosa* LYSZa5 could be found from BioProject No. PRJNA712958 and PRJNA712961 on NCBI.

## Results

### Hospital-acquired *P. aeruginosa* in the respiratory systems of COVID-19 patients

During routine screening of respiratory samples of COVID-19 patients, we discovered that two patients were colonized by *P. aeruginosa* with the same ST type. We have identified and isolated *P. aeruginosa* from 5 different critically ill patients in total. To investigate genetic adaptation and epidemiological link between these *P. aeruginosa* isolates, genomes of all isolates were sequences by Illumina HiSeq platform. Multi-Locus Sequence Typing (MLST) analysis indicated that four isolates collected from 2 patients are of the same type, *P. aeruginosa* ST1074, showing that these *P. aeruginosa* isolated from these two patients were hospital-acquired. We thus focused on these isolates to analyze their competitive advantage during colonization in COVID-19 environment. These four *P. aeruginosa* isolates were collected from respiratory samples of the two patients and named as LYSZa2, LYSZa3, LYSZa5 and LYSZa6 respectively. Among which, LYSZa2 and LYSZa3 were isolated from patient 1 with 3 days interval while LYSZa2 was isolated on day 12 of hospitalization. LYSZa5 and LYSZa6 were isolated from patient 2 with 15 days interval while LYSZa5 was isolated on day 17 of hospitalization. Genomes of the first isolate of each patient, LYSZa2 and LYSZa5, were further sequenced using Pacific Biosciences RSII sequencer for further analysis. We defined LYSZa2 and LYSZa5 as the ancestry isolates while LYSZa3 and LYSZa6 as the progeny isolates, and analyzed their general genomic characteristics.

### Characterization of LYSZa2 and LYSZa5 genomes, phylogeny and antibiotic resistance

Circular genomes of LYSZa2 and LYSZa5 were assembled with genome sizes of 6 638 980 bp and 6 638 990 bp, with 66.2% of GC respectively. Rearrangements were checked and manually curated after pairwise comparison of the two genomes. As predicted by PGAP, LYSZa2 genome contains 6183 genes among which 6046 features encode for proteins and 86 encode for RNAs, while LYSZa5 genome contains 6183 genes among which 6045 features encode for proteins and 86 encodes for RNAs.

Both of the genomes contain the same antimicrobial resistance genes predicted by ResFinder, including *crpP, aph(3’)-IIb, catB7, blaOXA-50, blaPAO*, and *fosA* against Fluoroquinolone, Aminoglycoside, Phenicol, Beta-lactam and Fosfomycin drugs. No difference in antimicrobial genes was identified between the ancestry and the progeny isolates. *In vitro* antimicrobial resistance of the isolates were tested using antibiotics including ceftazidime, piperacillin, cefoperazone/sulbactam, imipenem, aztreonam and levofloxacin. Obvious differences in resistance were observed between the ancestry and the progeny isolates. LYSZa5 is sensitive to imipenem, aztreonam and levofloxacin. However, LYSZa6 becomes resistance to imipenem, aztreonam and intermediate to levofloxacin after 15 days of evolution (Table S1). No obvious change in resistance to these drugs was observed between LYSZa2 and LYSZa3 probably due to the short evolving time (Table S1).

Genomic islands (GIs) on LYSZa2 and LYSZa5 genomes were predicted by IslandViewer4 (Table S2&S3). In total, 35 GIs on LYSZa2 and 36 GIs on LYSZa5 were predicted by at least one prediction method. Genomes of LYSZa2 and LYSZa5 were compared with genomes of five other *P. aeruginosa* strains including PAO1 reference strain and virulence strains including PA14, LESB58, SCV20265 and VFRPA04 (Figure 1A). Most of the GIs predicted are specific to LYSZa2 and LYSZa5 genomes (Figure 1A). Genes in these GIs are involved in transcriptional regulation, DNA restriction-modification, DNA repair, toxin-antitoxin and secretion systems, showing that these GIs are essential for *P. aeruginosa* survival and virulence during coinfection with SARS-CoV-2.

**Figure 1.**
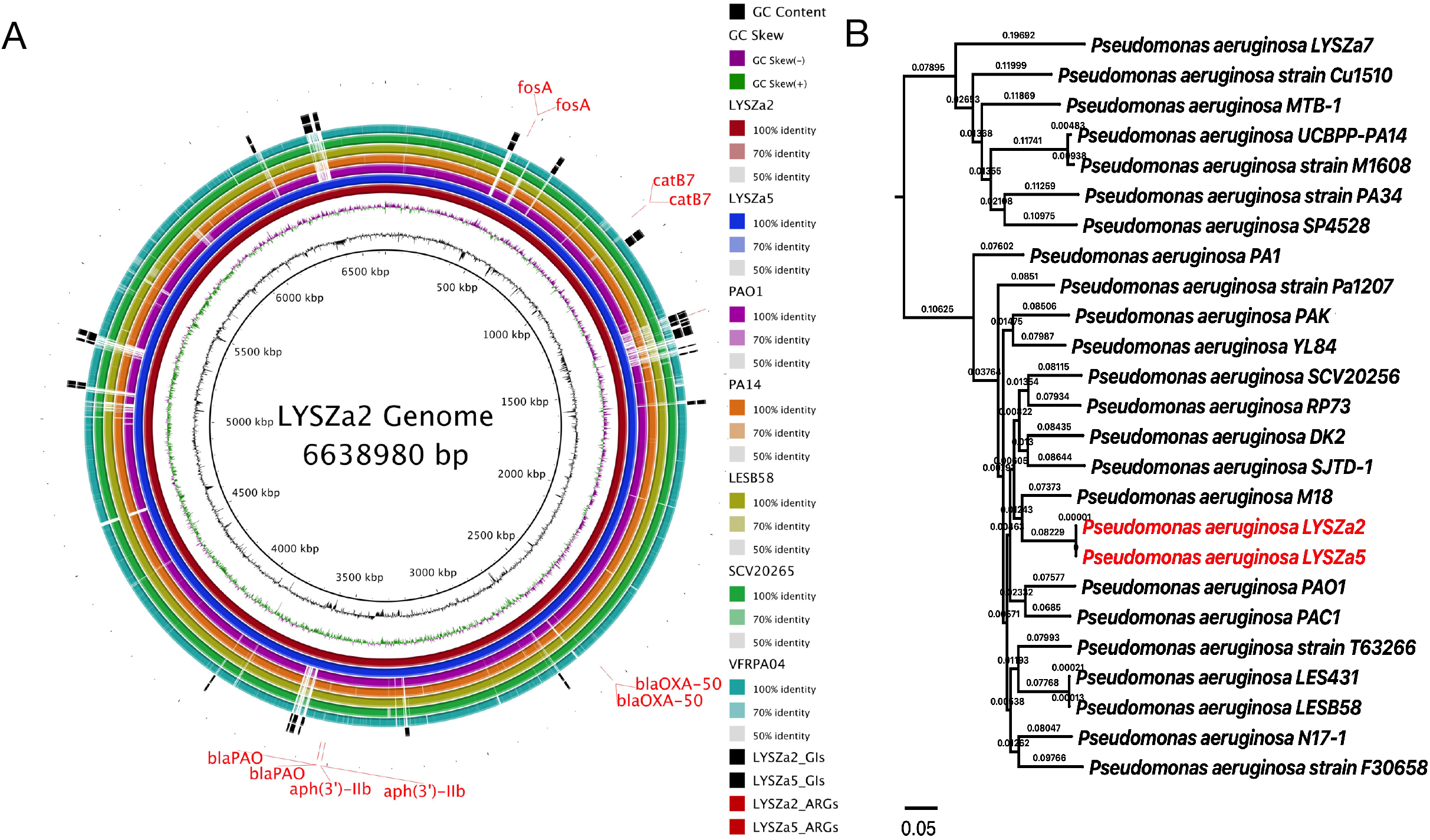
Ring Plot and Phylogenetic tree of LYSZa2 and LYSZa5. **(A)**Ring plot of LYSZa2 and LYSZa5 comparing with other *P. aeruginosa* genomes. From the innermost, ring 1: GC content (black); ring 2: GC skew (purple/green); ring 3: LYSZa2(dark red); ring 4: LYSZa5(navy); ring 5: PAO1 lab reference strain (violet); ring 6: PA14 clinical virulent strain (orange); ring 7: LESB58 hypervirulent clinical strain from CT patient (olive); ring 8: SCV20265 small colony variant strain (light green); ring 9: VRFPA04 multidrug resistant strain from keratitis patient(cyan); ring 10&11: GIs on LYSZa2 and LYSZa5 (black); ring 12&13: ARGs on LYSZa2 and LYSZa5 (red); **(B)** phylogenetic tree constructed using LYSZa2 and LYSZa5 genomes and 23 other *P. aeruginosa* genomes based on core genome alignment, LYSZa2 and LYSZa5 are highlighted in red.

We then traced the origin of these isolates by constructing phylogenetic tree using genomes of LYSZa2 and LYSZa5 with 22 other clinical or environmental *P. aeruginosa* genomes selected from NCBI/Pseudomonas genome database (Table S4) and one SARS-CoV-2 coinfecting strain published by our group recently, *P. aeruginosa* LYSZa7[33]. As seen from the phylogenetic tree (Figure 1B), LYSZa2 and LYSZa5 are closed related without evolutionary distance between them, and are in the same phylogenetic cluster with PAO1 reference strain and the hypervirulent isolate LESB58 from CF patient. Interestingly, great evolutionary distance was observed between these two isolates with *P. aeruginosa* LYSZa7, indicating the distinct evolutionary traits evolved between different *P. aeruginosa* strains in COVID-19 patients for adaptation.

Single nucleotide polymorphism (SNP) and other genome modifying events were assessed between the ancestry isolates and the progeny isolates respectively using PAO1 as reference (Table S5). 90 genomic modifying events including SNPs, insertion, deletion and replacement (Table S5) were identified in LYSZa3 with a dN/dS ratio of 0.875, indicating a negative selection during evolution, probably due to the short evolving time between these two isolates. 93 of such genomic modifying events (Table S5) were identified in LYSZa6 comparing to LYSZa5 with a dN/dS ratio of 1.114 indicating a positive selection. Common mutations were found between LYSZa3 and LYSZa6 on genes related to Type VI secretion system and iron transport (Table 1). Such observation indicated that *P. aeruginosa* undergoes both adaptive evolution and convergent evolution in COVID-19 patients to survive and modulate its virulence during coinfection. We then performed RNA sequencing to assess the changes at transcriptional level during evolution.

**Table 1.**
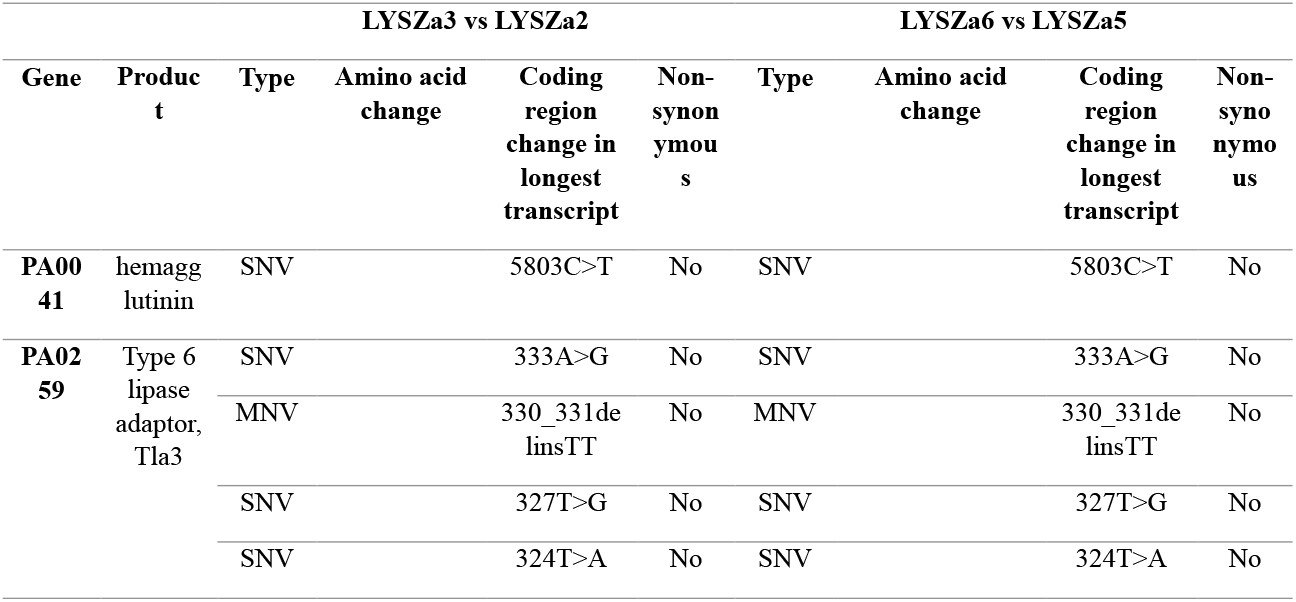
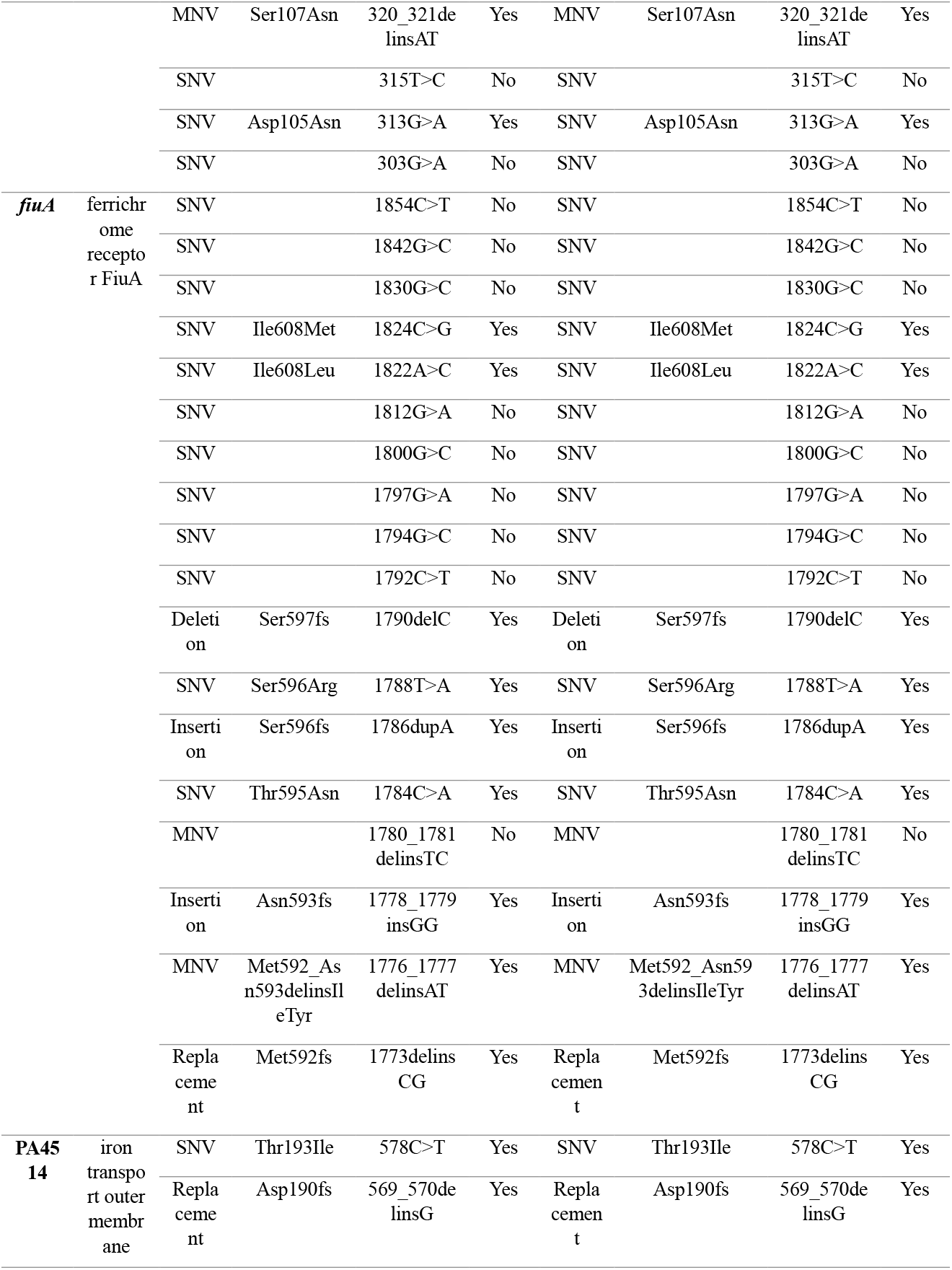
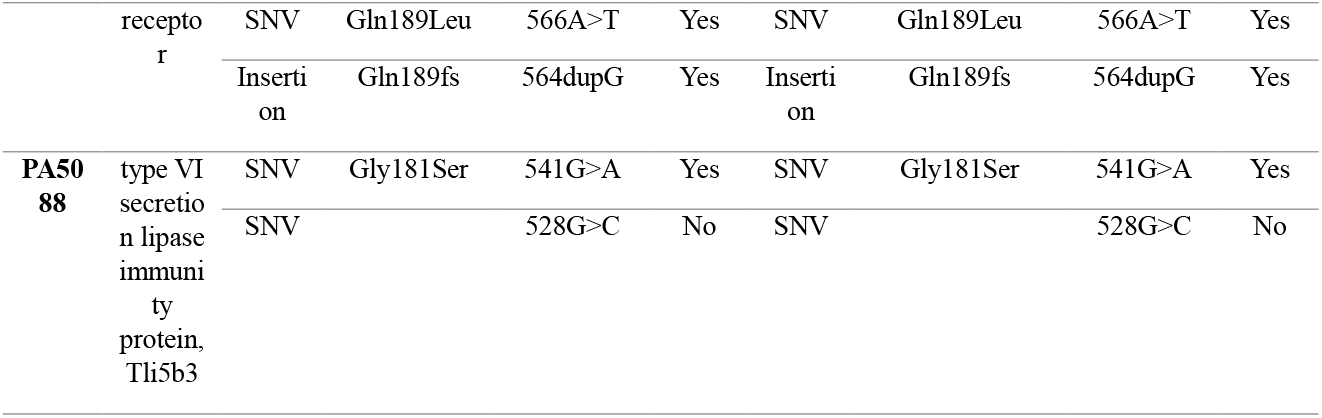
Common genetic mutations identified between LYSZa3 and LYSZa6. Full list of genetic mutations is included in Table S5.

### Alginate overproduction and attenuated expression of T6SS during evolution

To discover the transcriptional changes during adaptive evolution, we performed RNA sequencing of the isolates on Illumina HiSeq platform and analyzed differential gene expression between the ancestry isolates and the progeny isolates longitudinally. Moreover, we also compared the gene expression in parallel between the progeny cells from the two patients to learn the convergent changes of *P. aeruginosa* in different hosts. Based on the filtering criteria of fold change ≥ 4, adjusted p-value<0.05 and base mean ≥ 20, the expression of 129 genes were differentially regulated in LYSZa3 comparing to LYSZa2, among which 75 were upregulated and others were downregulated (Table S6). In LYSZa6 as comparing to LYSZa5, 242 genes were differentially expressed, among which 114 were upregulated and others were downregulated (Table S7). These differentially expressed genes are illustrated by heatmaps (Figure 2A&B), which display distinctive transcriptomic profiles of the progeny isolates comparing to the ancestry isolates. We further clustered the isolates using Principal Coordinates Analysis (PCoA) based on Bray Curtis dissimilarity to check the differences between isolates. As seen from PCoA plots (Figure 2C&D), LYSZa2 group and LYSZa5 group separated clearly from LYSZa3 group and LYSZa6 group respectively, indicating the ancestry isolates and the progeny isolates possessing distinct physiology.

**Figure 2.**
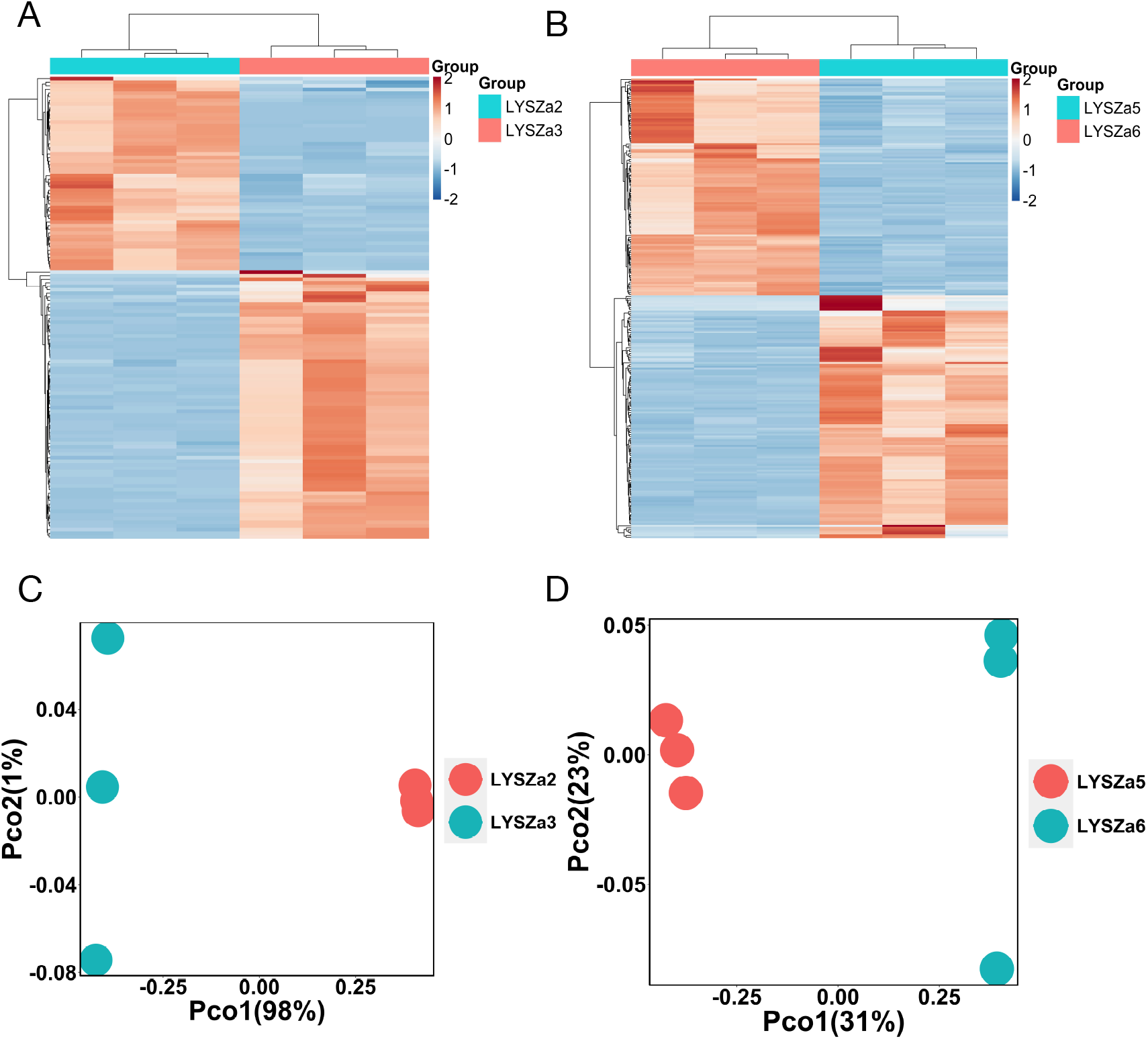
Heatmaps and PCoA plots of DEGs between the ancestry isolates and the progeny isolates. **(A)** heatmap of significant DEGs in LYSZa3 comparing to LYSZa2; **(B)** heatmap of significant DEGs in LYSZa6 comparing to LYSZa5; **(C)** PCoA plots of LYSZa3 group and LYSZa2 group based on DEGs using Bray Curtis dissimilarity; **(D)** PCoA plots of LYSZa6 group and LYSZa5 group based on DEGs using Bray Curtis dissimilarity.

We then performed Gene Ontology (GO) enrichment analysis using the differentially expressed genes (DEGs) to identify highly enriched functions in the progeny isolates comparing to the ancestry isolates (Figure 3). 11 GO consisting of 7 biological processes, 3 molecular functions and 1 cellular component were enriched in LYSZa3 comparing to LYSZa2 (Figure 3A). Whereas in LYSZa6 comparing to LYSZa5, 13 GO consisting of 7 biological processes, 5 molecular functions and 1 cellular component were highly enriched (Figure 3C). Among all of these enriched functions in both the progeny isolates, most significantly enriched biological processes are alginic acid biosynthetic process and protein secretion by the type VI secretion system (T6SS). 12 genes assigned to alginic acid biosynthetic process (*algD, algX, algA, algE, algF, algL, alg44, algJ, algK, alg8, algI, algG*) were all upregulated in LYSZa3 for 16.89 to 1091.52 folds (Table 2). The expression of same genes also increased significantly in LYSZa6 for 4.45 to 138.79 folds (Table 3). Eight genes assigned to protein secretion by T6SS, PA1657 (*hsiB2*), PA1658 (*hsiC2*), PA1659 (*hsiF2*), PA1660(*hsiG2*), PA1661(*hsiH2*), PA1662(*clpV2*), PA1663(*sfa2*), PA1666(*lip2*), were all significantly downregulated in LYSZa3 for 4.46 to 6.47 folds (Table 2). In LYSZa6, beside the same eight genes mentioned above, 7 other genes assigned to protein secretion by T6SS were also downregulated, including PA1656 (*hsiA2*), PA1665 (*fha2*), PA1667 (*hsiJ2*), PA1668 (*dotU2*), PA1669 (*icmF2*), *stk1* and *stp1*, for 4.46 to 17.11 folds (Table 3). Besides these genes assigned to T6SS by GO enrichment analysis, the expression of several other genes involved in T6SS, *hcpB, lip3* (PA2364), and *dotU3* (PA2362), were also decreased in LYSZa3 (Table 2). While the expression of 15 other genes involved in T6SS significantly decreased in LYSZa6, including *clpV1*, PA3904(PAAR4), PA2702(*tse2*), PA2774(*tse4*), PA2775(*tsi4*), *vgrG1*, PA0093(*tse6*), PA3905(*tecT*), PA0082(*tssA1*), PA2703(*tsi2*), PA0094(*eagT6*), PA3484(*tse3*), PA5266(*vgrG6*), *hcpA* and *hcpB* (Table 3). *P. aeruginosa* carries three types of T6SS, HSI-I, HSI-II and HSI-III respectively. As illustrated in Figure 4, the downregulated genes in both progeny isolates, especially those in LYSZa6, are mostly involved in HSI-II gene cluster, with several others scattered on another two gene clusters. Higher no. of DEGs observed in LYSZa6 comparing to LYSZa3 may consequently due to the longer evolving time of LYSZa6 in the host. The changes in the gene abundance assigned to different GO functions are illustrated in Figure 3B&D. There was dramatic increase in the gene abundance for alginic acid biosynthetic process and significant decrease in the gene abundance for protein secretion by T6SS in the progeny isolates. Such observation conveyed excessive biosynthesis of alginate and suppression of T6SS in the progeny isolates during colonization. The progeny isolates produce excessive alginate and exhibit more mucoid phenotype which matches with the DEG results (Figure 5A). As alginate plays an important role in biofilm architecture and development[34], we thus quantified biofilm formation of the isolates by crystal violet staining. As seen from Figure 5B, aligned with gene expression, an increase in the biofilm formation were observed in LYSZa3 comparing to those of LYSZa2. Similarly, more biofilm formation was observed in LYSZa6 as compared to LYSZa5. RNA-seq analysis showed the adaptive changes in genes responsible for alginate biosynthesis and T6SS protein secretion in the progeny isolates.

**Table 2.**
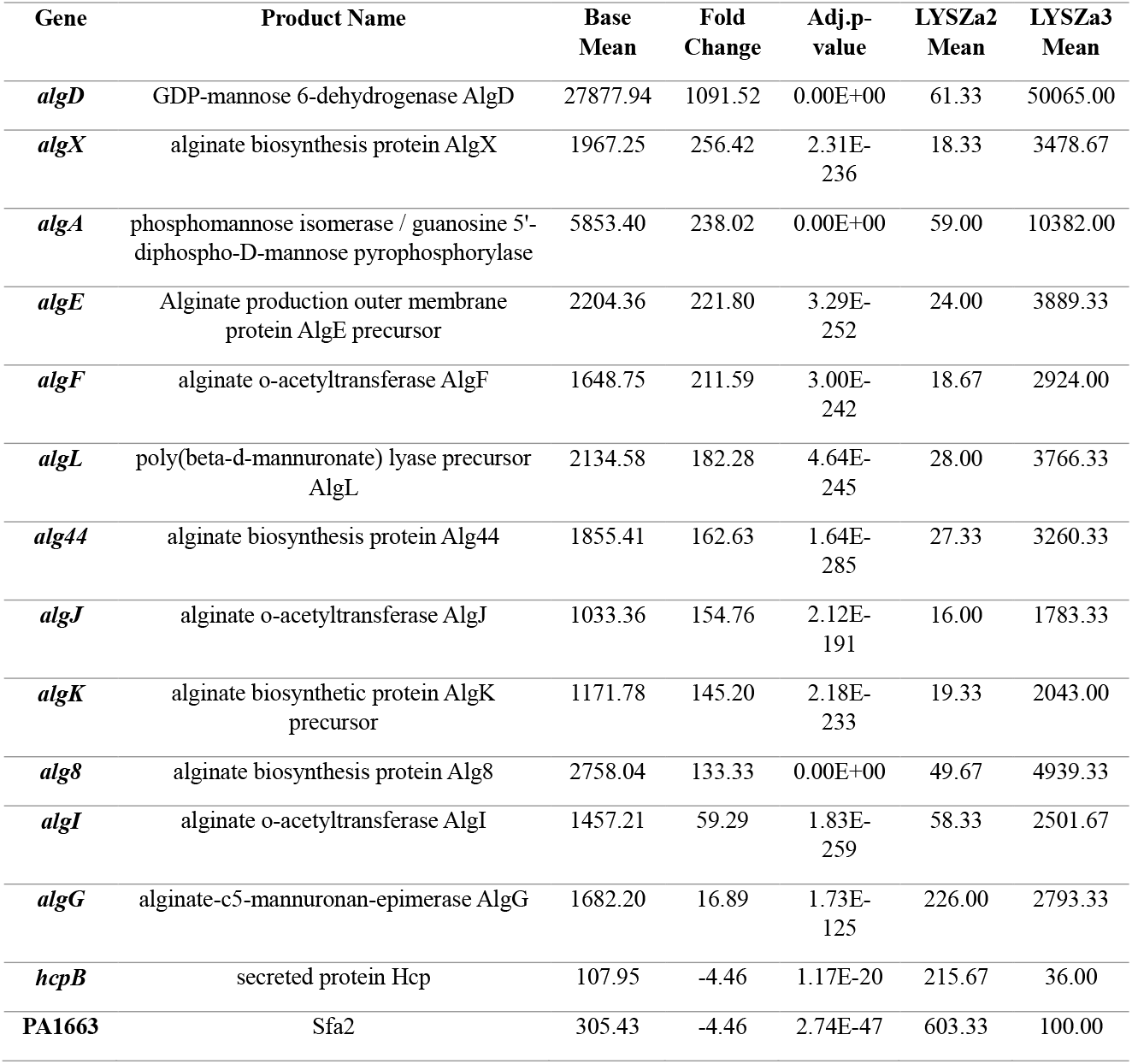
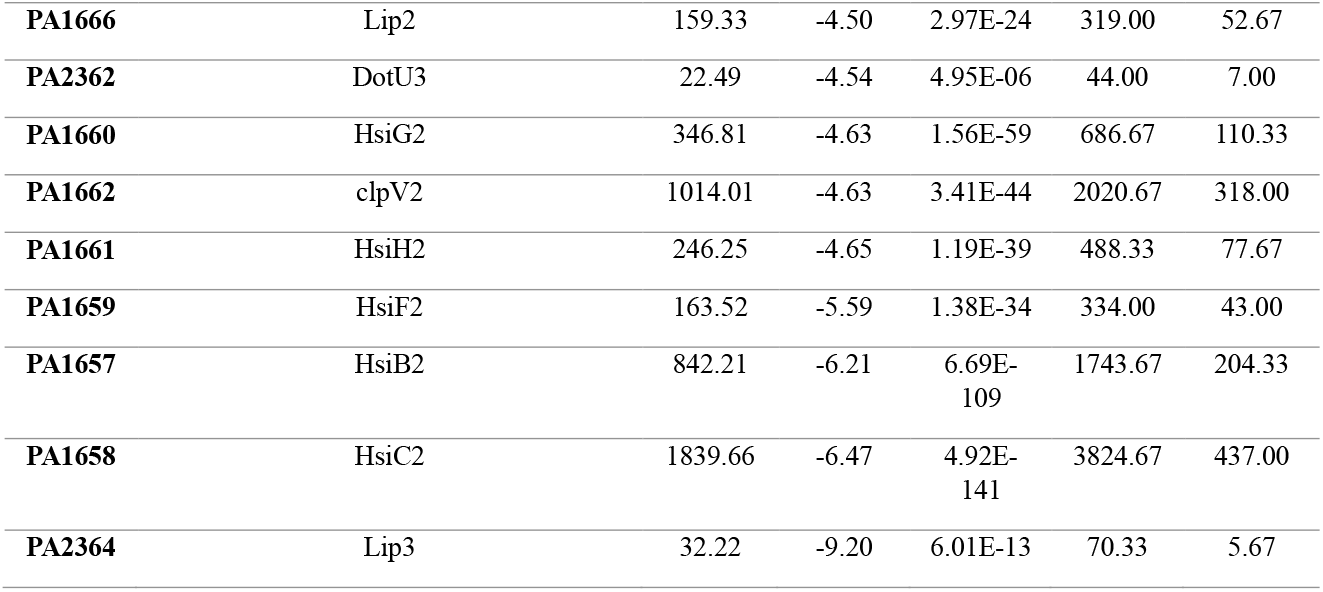
Selected DEGs in LYSZa3 comparing to LYSZa2 related to alginate biosynthesis and Type VI secretion system. Filtering criteria are fold change ≧4, adjusted p-value<0.05 and base mean ≧ 20. Full list of DEGs is included in Table S6.

| Gene | Product Name | Base Mean | Fold Change | Adj.p-value | LYSZa2 Mean | LYSZa3 Mean |
| --- | --- | --- | --- | --- | --- | --- |
| *algD* | GDP-mannose 6-dehydrogenase AlgD | 27877.94 | 1091.52 | 0.00E+00 | 61.33 | 50065.00 |
| *algX* | alginate biosynthesis protein AlgX | 1967.25 | 256.42 | 2.31E-236 | 18.33 | 3478.67 |
| *algA* | phosphomannose isomerase / guanosine 5'-diphospho-D-mannose pyrophosphorylase | 5853.40 | 238.02 | 0.00E+00 | 59.00 | 10382.00 |
| *algE* | Alginate production outer membrane protein AlgE precursor | 2204.36 | 221.80 | 3.29E-252 | 24.00 | 3889.33 |
| *algF* | alginate o-acetyltransferase AlgF | 1648.75 | 211.59 | 3.00E-242 | 18.67 | 2924.00 |
| *algL* | poly(beta-d-mannuronate) lyase precursor AlgL | 2134.58 | 182.28 | 4.64E-245 | 28.00 | 3766.33 |
| *alg44* | alginate biosynthesis protein Alg44 | 1855.41 | 162.63 | 1.64E-285 | 27.33 | 3260.33 |
| *algJ* | alginate o-acetyltransferase AlgJ | 1033.36 | 154.76 | 2.12E-191 | 16.00 | 1783.33 |
| *algK* | alginate biosynthetic protein AlgK precursor | 1171.78 | 145.20 | 2.18E-233 | 19.33 | 2043.00 |
| *alg8* | alginate biosynthesis protein Alg8 | 2758.04 | 133.33 | 0.00E+00 | 49.67 | 4939.33 |
| *algI* | alginate o-acetyltransferase AlgI | 1457.21 | 59.29 | 1.83E-259 | 58.33 | 2501.67 |
| *algG* | alginate-c5-mannuronan-epimerase AlgG | 1682.20 | 16.89 | 1.73E-125 | 226.00 | 2793.33 |
| *hcpB* | secreted protein Hcp | 107.95 | -4.46 | 1.17E-20 | 215.67 | 36.00 |
| **PA1663** | Sfa2 | 305.43 | -4.46 | 2.74E-47 | 603.33 | 100.00 |
| **PA1666** | Lip2 | 159.33 | -4.50 | 2.97E-24 | 319.00 | 52.67 |
| **PA2362** | DotU3 | 22.49 | -4.54 | 4.95E-06 | 44.00 | 7.00 |
| **PA1660** | HsiG2 | 346.81 | -4.63 | 1.56E-59 | 686.67 | 110.33 |
| **PA1662** | clpV2 | 1014.01 | -4.63 | 3.41E-44 | 2020.67 | 318.00 |
| **PA1661** | HsiH2 | 246.25 | -4.65 | 1.19E-39 | 488.33 | 77.67 |
| **PA1659** | HsiF2 | 163.52 | -5.59 | 1.38E-34 | 334.00 | 43.00 |
| **PA1657** | HsiB2 | 842.21 | -6.21 | 6.69E-109 | 1743.67 | 204.33 |
| **PA1658** | HsiC2 | 1839.66 | -6.47 | 4.92E-141 | 3824.67 | 437.00 |
| **PA2364** | Lip3 | 32.22 | -9.20 | 6.01E-13 | 70.33 | 5.67 |

**Table 3.** Selected DEGs in LYSZa6 comparing to LYSZa5 related to alginate biosynthesis and Type VI secretion system. Filtering criteria are fold change ≧4, adjusted p-value<0.05 and base mean ≧ 20. Full list of DEGs is included in Table S7.

| Gene | Product Name | Base Mean | Fold Change | Adj. p-value | LYSZa2 Mean | LYSZa3 Mean |
| --- | --- | --- | --- | --- | --- | --- |
| *algD* | GDP-mannose 6-dehydrogenase AlgD | 2956.90 | 138.79 | 2.47E-170 | 39.33 | 6375.00 |
| *algX* | alginate biosynthesis protein AlgX | 271.80 | 45.97 | 1.05E-75 | 10.67 | 585.67 |
| *algL* | poly(beta-d-mannuronate) lyase precursor AlgL | 283.61 | 44.85 | 1.98E-74 | 11.33 | 609.00 |
| *algJ* | alginate o-acetyltransferase AlgJ | 118.03 | 32.97 | 3.93E-33 | 6.33 | 248.67 |
| *algE* | Alginate production outer membrane protein AlgE precursor | 254.48 | 25.19 | 1.11E-61 | 18.33 | 535.33 |
| *algA* | phosphomannose isomerase / guanosine 5'-diphospho-D-mannose pyrophosphorylase | 481.88 | 21.52 | 9.22E-64 | 39.00 | 1000.67 |
| *algF* | alginate o-acetyltransferase AlgF | 152.00 | 18.68 | 7.50E-42 | 14.33 | 314.67 |
| *alg8* | alginate biosynthesis protein Alg8 | 292.51 | 14.91 | 1.09E-59 | 34.67 | 599.33 |
| *alg44* | alginate biosynthesis protein Alg44 | 195.56 | 14.43 | 3.38E-61 | 23.67 | 403.00 |
| *algK* | alginate biosynthetic protein AlgK precursor | 135.65 | 10.35 | 6.37E-39 | 21.67 | 272.67 |
| *algI* | alginate o-acetyltransferase AlgI | 218.14 | 8.40 | 7.47E-47 | 43.00 | 427.33 |
| *algG* | alginate-c5-mannuronan-epimerase AlgG | 356.62 | 4.45 | 4.15E-53 | 121.33 | 644.33 |
| *clpV1* | ClpV1 | 5315.67 | -4.08 | 7.48E-33 | 7854.33 | 2309.33 |
| **PA3904** | PAAR4 | 1457.06 | -4.17 | 3.79E-33 | 2178.00 | 631.67 |
| **PA2702** | Tse2 | 280.90 | -4.35 | 2.50E-29 | 419.33 | 117.33 |
| **PA2774** | Tse4 | 296.61 | -4.37 | 4.85E-26 | 461.67 | 122.33 |
| **PA2775** | Tsi4 | 133.81 | -4.71 | 7.24E-19 | 212.67 | 52.33 |
| *vgrG1* | VgrG1 | 2637.35 | -4.74 | 1.56E-50 | 3969.33 | 1016.67 |
| **PA0093** | Tse6 | 947.79 | -4.91 | 3.88E-46 | 1469.00 | 360.00 |
| **PA3905** | type VI effector chaperone for Tox-Rease,<br>TecT | 1073.56 | -4.95 | 1.10E-53 | 1671.00 | 403.00 |
| **PA0082** | TssA1 | 1449.46 | -5.32 | 1.76E-55 | 2265.00 | 515.00 |
| **PA2703** | Tsi2 | 98.26 | -5.56 | 5.99E-24 | 156.67 | 33.00 |
| **PA0094** | EagT6 | 251.06 | -5.79 | 1.43E-38 | 401.67 | 82.00 |
| **PA3484** | Tse3 | 544.56 | -5.86 | 1.50E-54 | 869.33 | 174.33 |
| *stk1* | Stk1 | 263.85 | -5.90 | 1.38E-56 | 423.00 | 84.67 |
| **PA5266** | VgrG6 | 396.48 | -6.81 | 6.50E-47 | 654.33 | 113.33 |
| **PA1667** | HsiJ2 | 678.54 | -6.99 | 2.41E-100 | 1119.67 | 188.67 |
| **PA1656** | HsiA2 | 3257.03 | -7.01 | 4.84E-80 | 5349.67 | 911.33 |
| **PA1662** | clpV2 | 2538.08 | -8.98 | 3.88E-108 | 4332.33 | 564.33 |
| **PA1661** | HsiH2 | 693.51 | -9.91 | 1.55E-110 | 1186.67 | 142.00 |
| **PA1666** | Lip2 | 340.97 | -10.48 | 2.48E-72 | 589.00 | 65.67 |
| *stp1* | Stp1 | 327.95 | -10.61 | 1.67E-70 | 562.67 | 62.67 |
| *hcpA* | secreted protein Hcp | 72.34 | -10.75 | 7.25E-26 | 125.33 | 13.67 |
| **PA1665** | Fha2 | 840.73 | -10.96 | 1.12E-108 | 1456.67 | 155.33 |
| **PA1660** | HsiG2 | 907.03 | -11.05 | 1.78E-125 | 1551.67 | 167.33 |
| **PA1669** | IcmF2 | 2052.74 | -11.08 | 6.20E-107 | 3547.33 | 375.67 |
| *hcpB* | secreted protein Hcp | 204.27 | -11.30 | 8.81E-47 | 358.00 | 36.67 |
| **PA1668** | DotU2 | 487.58 | -11.30 | 1.39E-105 | 836.33 | 88.00 |
| **PA1663** | Sfa2 | 883.17 | -12.92 | 4.47E-74 | 1556.67 | 139.33 |
| **PA3334** | Acp3 | 236.04 | -13.79 | 1.82E-73 | 415.67 | 35.33 |
| **PA1657** | HsiB2 | 2539.11 | -14.39 | 1.32E-112 | 4445.00 | 366.00 |
| PA1659 | HsiF2 | 480.65 | -17.06 | 4.25E-89 | 843.33 | 59.00 |
| PA1658 | HsiC2 | 5746.91 | -17.11 | 4.82E-66 | 10179.00 | 695.67 |

**Figure 3.**
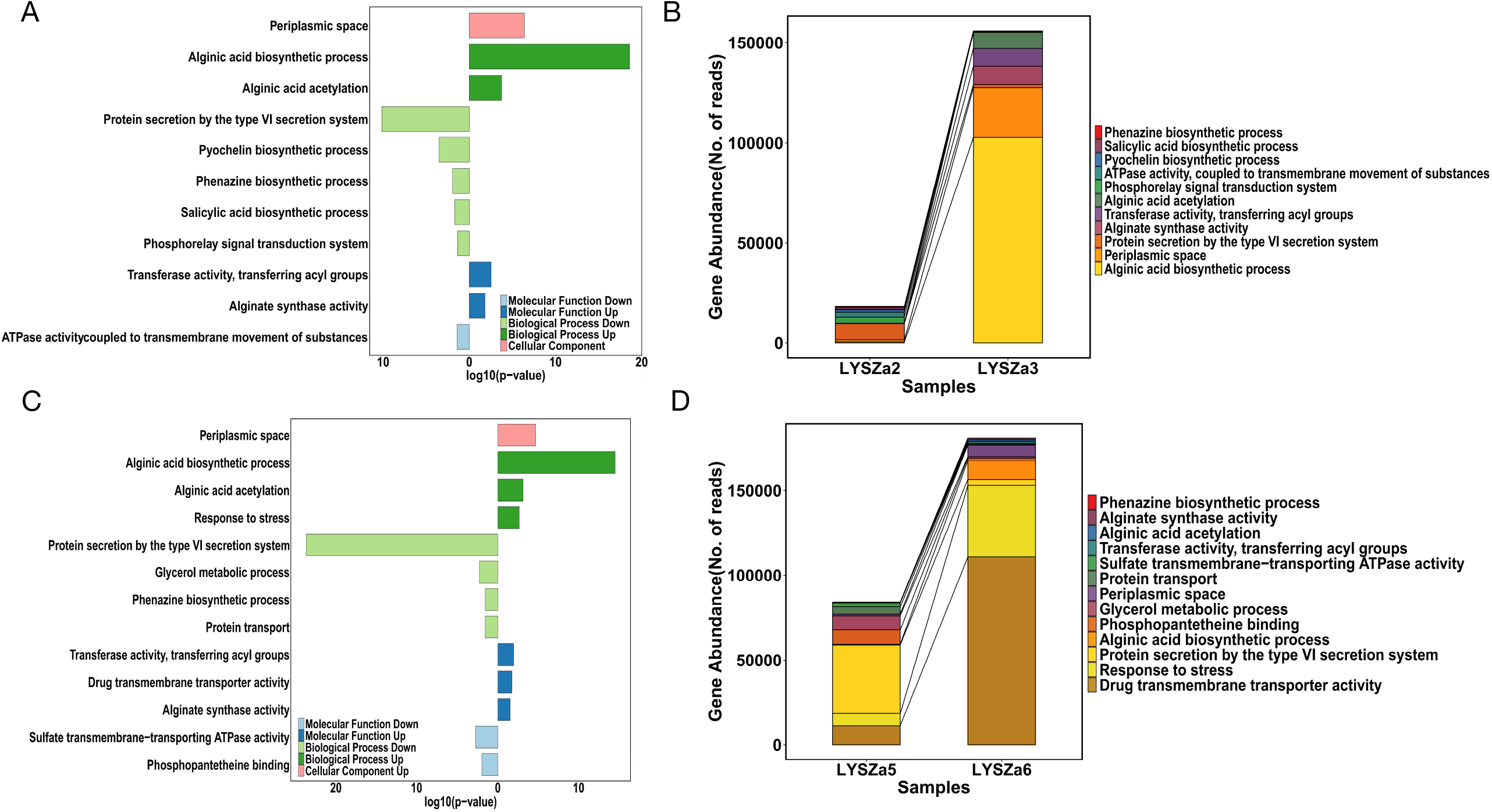
GO enrichment analysis and variation in gene abundance in each GO. **(A)** GO enriched in LYSZa3 comparing to LYSZa2, bars towards right-hand side showing genes assigned to the functions were upregulated, bars towards left-hand side showing genes assigned to the functions were downregulated, up- and down-regulation are indicated in the legend as well, p-value < 0.05; **(B)** Variation in gene abundance involved in the enriched GO functions listed in A in LYSZa2 and LYSZa3; **(C)** GO enriched in LYSZa6 comparing to LYSZa5, bars towards right-hand side showing genes assigned to the functions were upregulated, bars towards left-hand side showing genes assigned to the functions were downregulated, up- and down-regulation are indicated in the legend as well, p-value < 0.05; **(D)** Variation in gene abundance involved in the enriched GO functions listed in C in LYSZa5 and LYSZa6.

**Figure 4.**
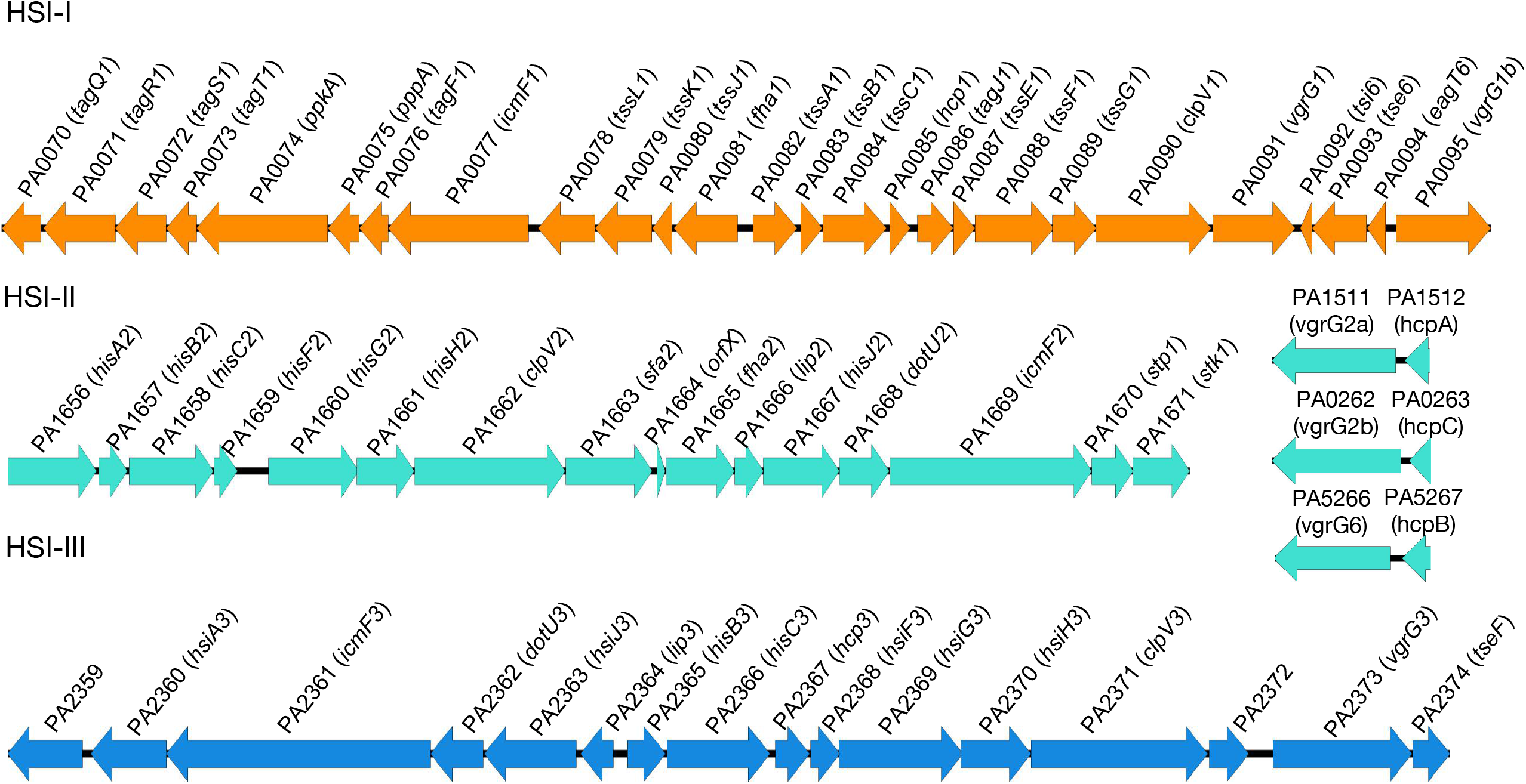
Schematic figure of HSI-I, HSI-II and HSI-III gene clusters. Genes downregulated in LYSZa3 are denoted by purple crosses; genes downregulated in LYSZa6 are denoted by red triangles. Gene name was obtained from Pseudomonas Genome Database [49].

**Figure 5.**
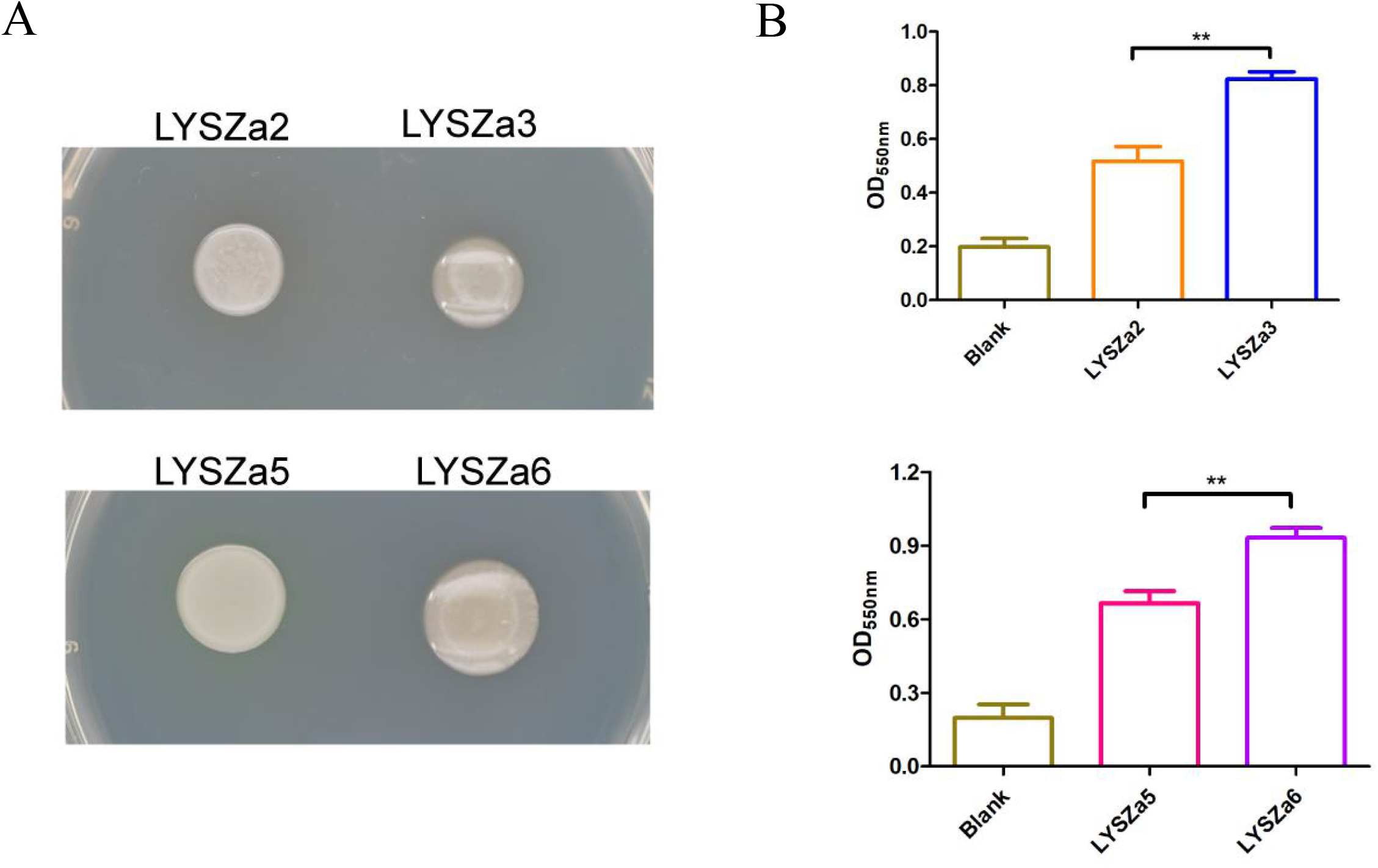
Phenotypic analysis. **(A)** Colony morphologies of the isolates; **(B)** Biofilm quantification tests by crystal violet staining, **: p-value < 0.05; Upper panel: biofilm quantification test of LYSZa2 and LYSZa3, lower panel: biofilm quantification test of LYSZa5 and LYSZa6.

In addition, 74 common genes in total were found to be differentially regulated in both progeny isolates, LYSZa3 and LYSZa6 (Table S8), indicating convergent gene regulation of these two isolates during colonization in different patients. Similar to the results observed from adaptive changes, most of the common genes with specific functions are involved in alginate biosynthesis and T6SS, which are the convergent changes in *P. aeruginosa* in different hosts during coinfection with SARS-CoV-2 virus. More genes were differentially regulated in LYSZa6 than LYSZa3 probably due to the longer evolving time between LYSZa5 and LYSZa6. We observed increased alginate production and elevated biofilm formation of the progeny isolates from the phenotypic assays. T6SS is used as a weapon by *P. aeruginosa* to compete with neighboring bacteria cells and to combat with the host cells in order to gain surviving advantage in the polymicrobial environment in the respiratory system. Suppression in T6SS would result in reduction in its advantage to outcompete other bacteria and its virulence to escape from host immune clearance. We thus performed bacterial competition test and cytotoxicity assay of the isolates to confirm the changes in the anti-prokaryotic and anti-eukaryotic capability between the ancestry isolates and the progeny isolates.

### Decrease in bactericidal activity of *P. aeruginosa* due to suppression of T6SS

To test the reduction in the competitiveness of the progeny isolates with bacterial neighbors due to suppression of T6SS, we examined the capability of the isolates in killing *E. coli* cells through bacterial competition assay. The isolates were mixed and cultured with *E. coli/placZ* in 1:1 ratio respectively. The mixtures were diluted to 10^−3^ while triplicate colonies of each dilution were then cultured on plate containing X-gal to test the killing efficiency (Figure 6A&B). *E. coli/pLacZ* digested X-gal and appeared as blue colonies. Survival of *E. coli* was determined both qualitatively by visualizing the intensity of blue pigment of the colonies and quantitatively by counting the blue *E. coli* CFUs left. As observed from Figure 6A&B, almost no trace of blue pigment could be observed from the mixture of the ancestry isolates, LYSZa2 and LYSZa5, with *E. coli*. More intensive blue colonies were observed from the mixed cultures of the progeny isolates, LYSZa3 and LYSZa6, with *E. coli*, indicating disadvantage of these isolates to outcompete *E. coli* comparing to their respective ancestry isolates. Same trend was also observed from quantitative tests illustrated in Figure 6C where higher numbers of *E. coli* CFU were obtained after coculturing with LYSZa3 and LYSZa6 comparing with those culturing with LYSZa2 and LYSZa5 respectively. Such results indicated a decrease in the bactericidal activity of the progeny isolates during bacterial competition consequently due to the inhibition of T6SS.

**Figure 6.**
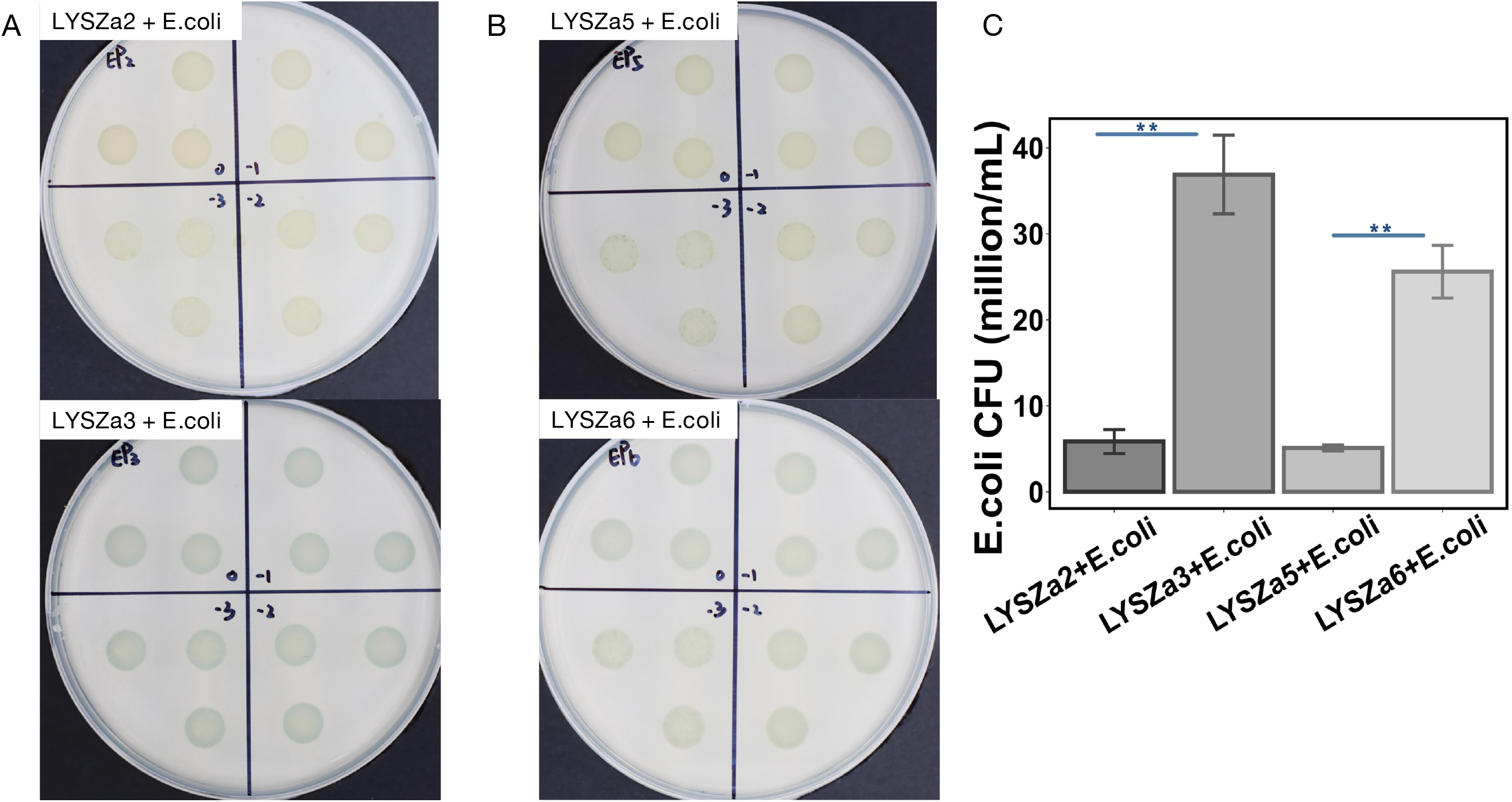
Bacterial competition assay between the isolates and *E. coli*. Blue pigment intensity indicates the survival of *E. coli*.**(A)** Upper panel: competition between LYSZa2 and *E. coli*; lower panel: competition between LYSZa3 and *E. coli*, number in each section denotes the dilution factor, ranged from 10^0^ to 10^−3^; **(B)** Upper panel: competition between LYSZa5 and *E. coli*; lower panel: competition between LYSZa6 and *E. coli*, number in each section denotes the dilution factor, ranged from 10^0^ to 10^−3^; **(C)** Counts of *E. coli* CFU left in each competition, **: p-value<0.05.

### Weakened macrophage cytotoxicity of *P. aeruginosa* after T6SS suppression

We then performed macrophage killing assay to assess the decrease in cytotoxicity of the progeny isolates to eukaryotic cells upon T6SS suppression. Cultures of the isolates were added to infect macrophages individually. Relative lactate dehydrogenase (LDH) release was measured to determine the death of macrophages. More macrophages are killed, higher LDH will be released. As illustrated in Figure 7, LDH released by macrophages infected by LYSZa3 was much lower than that of LYSZa2. Similar results were observed from LYSZa5 and LYSZa6 where macrophages infected by LYSZa5 released a significant higher level of LDH comparing to LYSZa6. LYSZa3 and LYSZa6 possess weaker cytotoxicity comparing to their ancestral isolates. Such results suggested that these *P. aeruginosa* isolates produced attenuated virulence towards their eukaryotic hosts due to the downregulation of T6SS genes after evolution.

**Figure 7.**
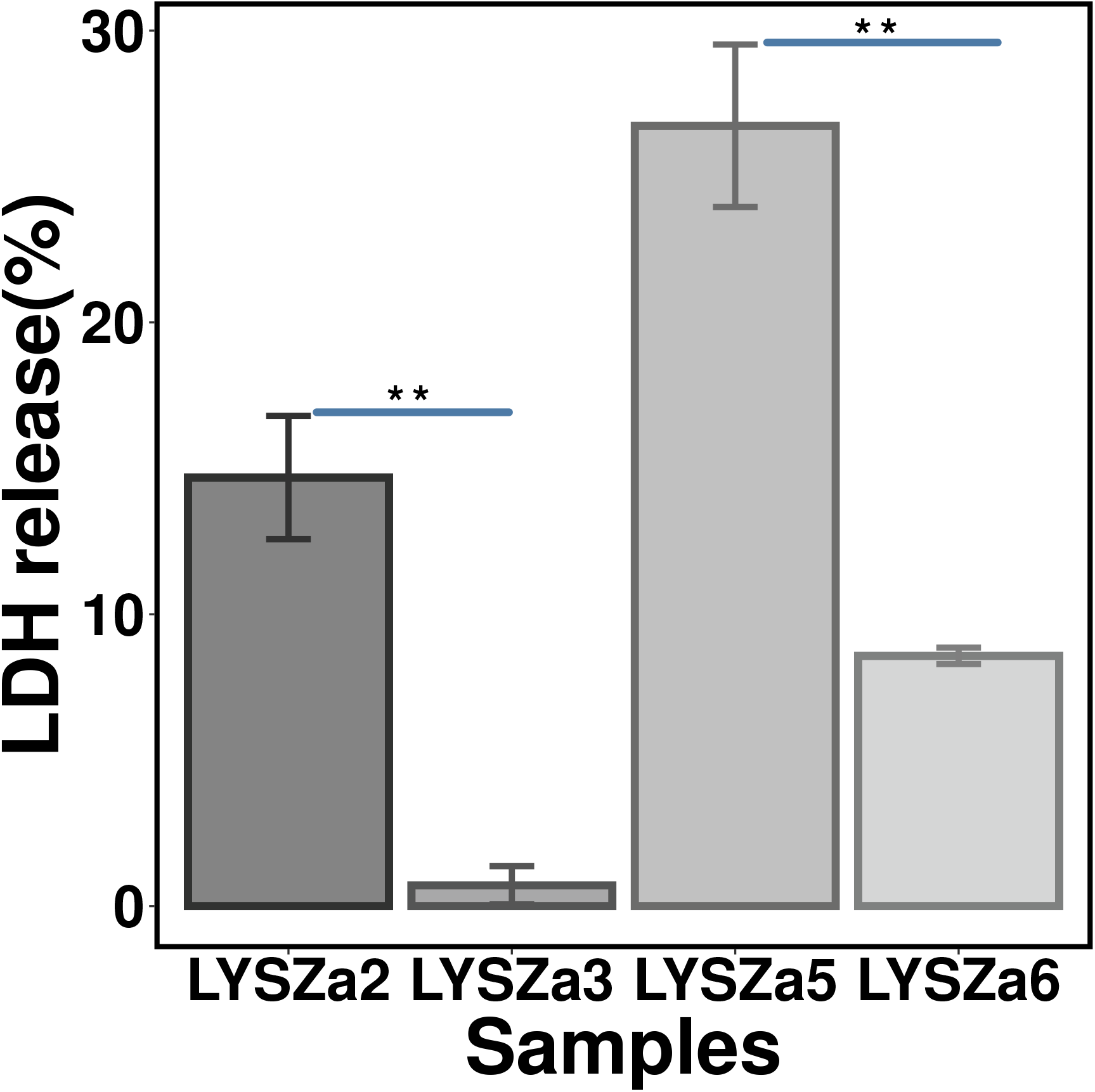
Macrophage cytotoxicity assay of the ancestry isolates and the progeny isolates. Killing of macrophage is illustrated by the percent of LDH released, **: p-value<0.05.

## Discussion

Upon the urgent need to prevent the spread of SARS-CoV-2 and equip people against the virus, clinical doctors and researchers put their attention on investigating the pathogenicity and epidemiology of the virus, and developing treatment measures and vaccines to combat it. However, the increased occurrence of illness and death due to bacterial coinfection and its complications indicated that there is a much underestimated and neglected influence of bacterial coinfection on the disease progression in COVID-19 patients. Here in this study, we isolated the ancestry and the progeny isolate pairs of hospital-acquired *P. aeruginosa* from two critically ill COVID-19 patients to investigate its adaptive and convergent evolution during coinfection with SARS-CoV-2 virus. We found that *P. aeruginosa* upregulates its alginate biosynthesis and downregulates its T6SS to increase its fitness in the niche and reduce its virulence to escape from host clearance for longer colonization. To the best of our knowledge, this is the first study describing the evolution of *P. aeruginosa* during coinfection with SARS-CoV-2 virus in COVID-19 patients.

The isolates collected from the two patients belongs to the same sequence type. In addition, the ancestry isolates appeared after more than 10 days of hospitalization, they are therefore hospital-acquired *P. aeruginosa* strain. Genomic sequencing analysis revealed that the genomes of the isolates carry specific GIs for DNA repair and protein secretion systems, and ARGs for multiple drug classes. Common genomic modification events identified between the progeny isolates and the ancestral isolates indicated the genomic changes in genes related to T6SS, *tla3* and *tli5b3*. However, no change was observed in these two genes at transcriptional level most probably due to the short evolving time and the complexity of gene regulation cascades. Although the expression of these two genes were not affected, we indeed observed lots of significant changes at transcriptional level.

Through differential gene expression analysis, we found that genes involved in alginate biosynthesis were upregulated greatly while genes involved in T6SS protein secretion system were significantly downregulated. Alginate is an essential EPS component converting non-mucoid *P. aeruginosa* to mucoid phenotype, and is important for biofilm structure and antimicrobial resistance [34]. Clinical *P. aeruginosa* strains overproducing alginate isolated from CF patients enhanced biofilm formation and interfered with host immune responses leading to the aggravation of disease conditions and poor prognosis [35, 36]. Moreover, excessive biosynthesis of alginate in *P. aeruginosa* promotes its coinfection with other pathogens such as *S. aureus* and *B. cenocepacia* and increases their persistence in CF infections [37, 38]. Overproduction of alginate and increased biofilm formation were observed in the progeny strains in this study, inferring the potential mechanism adopted by *P. aeruginosa* in COVID-19 patients to interfere with host defense systems. Thick alginate layer is also probably a key contributing factor to the increase in the antimicrobial resistance of the progeny isolate observed. Excessive alginate may also facilitate coexistence of *P. aeruginosa* and other coinfecting pathogens in COVID-19 for secondary bacterial infection. Further analysis is needed for deeper investigation on the role of alginate in the microbial community in COVID-19 patients.

A more notable observation in this study is the attenuation of T6SS in the progeny isolates. The expression of genes involved in T6SS protein secretion were significantly downregulated in both the progeny isolates. *P. aeruginosa* T6SS functions as a phage-like toxin delivery apparatus to transport toxic effectors into surrounding competing bacterial cells and also host cells to gain survival advantages and invade host cells [39]. *P. aeruginosa* carries three types of T6SS, being HSI-I, HSI-II, and HSI-III respectively. HSI-I is dedicated to inter-bacterial competition while HSI-II and HSI-III, which are less studied, associate with both anti-prokaryotic and anti-eukaryotic functions [40, 41]. Three gene clusters HSI-I, HSI-II and HSI-III contain *hcp, clpV, vgrG, tss/his* genes and other genes including *ppkA, pppA, tag, sfa, fha,lip, dotU, icmF, stp* and *stk* genes for the full function of T6SS apparatus (Figure 4)[39, 42]. In both the progeny isolates, especially in LYSZa6, the expression of most of HSI-II genes and associated genes were downregulated significantly (Figure 4). Such results indicated a general inhibition of HSI-II T6SS in the *P. aeruginosa* isolates during convergent evolution in the patients to adapt to COVID-19 environment. As HSI-II T6SS associates with anti-bacterial and anti-host functions, we tested both the bacterial killing capacity and macrophage cytotoxicity of the isolates. The results confirmed the decrease in the virulence to both neighbouring bacteria and host cells of the progeny isolates due to reduction in T6SS expression. Previous studies had demonstrated that *P. aeruginosa* abrogated its T6SS to adapt to the host environment by genetic modification on genes such as *vgrG* and *vgrG4* during chronic infection of cystic fibrosis [16, 43]. Here we showed that *P. aeruginosa* attenuated its T6SS at transcriptional level to gain fitness and survival advantages during acute infection of COVID-19 pneumonia. Moreover, attenuation of T6SS in *P. aeruginosa* may give opportunities to other bacterial pathogens to outcompete itself and making host cells susceptible to subsequent infections. A recent research showed that *P. aeruginosa* abrogating its T6SS could be outcompeted by coinfecting *B*.*cenocepaica* in CF patients in an age- and T6SS-dependent manner and predisposing host to superinfection of *B*.*cenocepaica* [44]. *P. aeruginosa* attenuating its T6SS in COVID-19 patients could probably play a pivotal role in coexisting with other pathogenic strains in the polymicrobial environment and promoting subsequent infection in COVID-19 patients. Further study is indeed needed to make a deeper investigation on the role of *P. aeruginosa* played in SARS-CoV-2 infected environment. In this study, we demonstrated that *P. aeruginosa* adapted to SARS-CoV-2 infected environment by adaptive and convergent evolution in alginate biosynthesis and T6SS gene expression for enhancing biofilm formation, increasing antimicrobial resistance and gaining fitness for survival and colonization during the acute COVID-19 infection. This study will contribute to the understanding of adaptation and convergent evolution of *P. aeruginosa* which in turn helps in the precise treatment decision and novel drug target discovery to eliminate *P. aeruginosa* coinfection in COVID-19 patients.

## Acknowledgements

This work was supported by Guangdong Natural Science Foundation for Distinguished Young Scholar[2020B1515020003] and Start-up Grants [Y01416106 and Y01416206] from the Southern University of Science and Technology (SUSTech) to Dr. Liang Yang, Science and Technology Program of Shenzhen [JCYJ20190809144005609] and Guangdong Basic and Applied Basic Research Foundation [2020A1515010586] to Dr. Jiuxin Qu, and Grant from Bill & Melinda Gates Foundation to Dr. Lei Liu. The authors declare that they have no competing interests.

